# Improved prediction of chronological age from DNA methylation limits it as a biomarker of ageing

**DOI:** 10.1101/327890

**Authors:** Qian Zhang, Costanza L. Vallerga, Rosie M Walker, Tian Lin, Anjali K. Henders, Grant W. Montgomery, Ji He, Dongsheng Fan, Javed Fowdar, Martin Kennedy, Toni Pitcher, John Pearson, Glenda Halliday, John B. Kwok, Ian Hickie, Simon Lewis, Tim Anderson, Peter A. Silburn, George D. Mellick, Sarah E. Harris, Paul Redmond, Alison D. Murray, David J. Porteous, Christopher S. Haley, Kathryn L. Evans, Andrew M. McIntosh, Jian Yang, Jacob Gratten, Riccardo E. Marioni, Naomi R. Wray, Ian J. Deary, Allan F. McRae, Peter M. Visscher

## Abstract

DNA methylation is associated with age. The deviation of age predicted from DNA methylation from actual age has been proposed as a biomarker for ageing. However, a better prediction of chronological age implies less opportunity for biological age. Here we used 13,661 samples (from blood and saliva) in the age range of 2 to 104 years from 14 cohorts measured on Illumina HumanMethylation450/EPIC arrays to perform prediction analyses. We show that increasing the sample size achieves a smaller prediction error and higher correlations in test datasets. We demonstrate that smaller prediction errors provide a limit to how much variation in biological ageing can be captured by methylation and provide evidence that age predictors from small samples are prone to confounding by cell composition. Our predictor shows a similar or better performance in non-blood tissues including saliva, endometrium, breast, liver, adipose and muscle, compared with Horvath’s across-tissue age predictor.

## Introduction

Ageing as a complex biological phenomenon is related to diseases and mortality^1,2^, and chronological age has been widely used as a marker of ageing due to ease and accuracy ofmeasurement^1^. However, chronological age is not necessarily a good predictor of biological ageing since individuals with the same chronological age can vary in health, especially in later life^3^. Therefore, biomarkers of ageing have become popular as they can indicate the presence or severity of some disease states^4,5^. In 2013, Hannum *et al*. and Horvath built age predictors based on DNA methylation and implemented them as biomarkers of ageing^6,7^. DNA methylation as a part of the epigenome plays an essential role in the regulation of gene expression in the human body^8,9^. Unlike DNA which is (mostly) stable across the lifetime of an individual, DNA methylation is dynamic, and previous studies have discovered a number of CpG sites associated with chronological age^10-12^. The age predictor developed by Hannum et al. was based on 482 blood samples with methylation measured on the Illumina 450K methylation arrays, and they reported a correlation of 0.91 and a Root Mean Square Error (RMSE) of 4.9 years in their test set^6^. Horvath’s age predictor was based on 8,000 samples from different tissues and cell types, and probes of these samples were from the Illumina 27K DNA methylation arrays. He reported a correlation of 0.96 and a Median Absolute Deviation (MAD) of 3.6 years in the test set. Age Acceleration Residuals (AAR) is defined as the residuals from regressing predicted age on chronological age. It has been reported to be associated with mortality, obesity and other complex traits^13-16^.

According to its definition, AAR is the prediction error in a chronological age predictor. Although previous survival analysis showed a significant association between AAR and mortality^13^, AAR was found not to be a mitotic clock^17^. Therefore, whether the significance of that association is inflated (e.g. by potential confounders in the error) needs to be investigated. To study the use of predicted age from DNA methylation as a biomarker of ageing, we calculated AAR based on age predictors with different prediction accuracy and investigated the relationship between prediction accuracy and the significance of AAR in survival analysis. We also investigated the effect of training sample size and statistical method on age prediction.

In the present study, we built DNA methylation-based age predictors by integrating 13,661 samples (13,402 from blood and 259 from saliva) measured on 450K DNA methylation arrays and Illumina EPIC (850K) arrays. Two approaches were evaluated: Elastic Net ^18^ and Best Linear Unbiased Prediction (BLUP)^19^. We discussed the implications of our results for the scope and utility of DNA methylation based age predictor as a biomarker for biological ageing. We also explored the factors that explain prediction accuracy. Finally, the performance of predictors on samples from tissues other than blood was investigated.

## Results

### Availability of DNA methylation in age prediction

We downloaded eight datasets from the public domain and used six datasets from our own studies **(Table 1)**. All data underwent identical quality control criteria before statistical analyses **(Material and Methods)**.

**Table 1:** Description of DNA methylation cohorts.

| COHORT<sup>1</sup> | SAMPLE SIZE<sup>2</sup> | NUMBER OF SAMPLES WITH VALID AGE | MEAN AGE(SD) | AGE RANGE | SOURCE | DISEASE |
| --- | --- | --- | --- | --- | --- | --- |
| **LBC1921**<sup>32,33</sup> | 692 | 692 | 82.3 (4.3) | [77.8, 90.6] | blood | Not Available |
| **LBC1936**<sup>32,33</sup> | 2326 | 2326 | 72.4 (2.8) | [67.7, 77.7] | blood | Not Available |
| **BSGS**<sup>31</sup> | 614 | 614 | 21.4 (14.1) | [9.9, 74.9] | blood | Not Available |
| **SGPD** | 1962 | 1556 | 67.2 (9.5) | [23.0,104.0] | blood | Parkinson's Disease: 988, Control: 974 |
| **MND**<sup>36</sup> | 695 | 600 | 45.2 (15.0) | [17.0,76.0] | blood | Motor Neuron Disease (MND): 497, Control: 198 |
| **GS**<sup>34,35</sup> | 5101 | 5100 | 48.5 (14.0) | [18.0,94.5] | blood | Not Available |
| **GSE72775**<sup>51</sup> | 335 | 335 | 70.2 (10.3) | [36.5,90.5] | blood | Not Available |
| **GSE78874**<sup>51</sup> | 259 | 259 | 68.8(9.7) | [36.0,88.0] | saliva | Not Available |
| **GSE72773**<sup>51</sup> | 310 | 310 | 65.6 (13.9) | [35.1,91.9] | blood | Not Available |
| **GSE72777**<sup>51</sup> | 46 | 46 | 14.7 (10.4) | [2.2,35.0] | blood | Not Available |
| **GSE41169**<sup>52</sup> | 95 | 95 | 31.6 (10.3) | [18.0,65.0] | blood | Schizophrenia:62, Control:33 |
| **GSE40279**<sup>6</sup> | 656 | 656 | 64.0 (14.7) | [19.0,101.0] | blood | Not Available |
| **GSE42861**<sup>53</sup> | 689 | 689 | 51.9 (11.8) | [18.0,70.0] | blood | Rheumatoid Arthritis:354, Control:335 |
| **GSE53740**<sup>54</sup> | 384 | 383 | 67.8(9.6) | [34.0,93.0] | blood | Alzheimer's Disease:15, Corticobasal Degeneration:1, Frontotemporal Dementia (FTD):121, FTD/MND:7, Progressive Supranuclear Palsy:43, Control:193, Unknown:4 |

<sup>1</sup> LBC = Lothian Birth Cohort; BSGS = Brisbane Systems Genomics Study; SGPD = Systems Genomic of Parkinson's Disease consortium; MND = Motor Neuron Disease cohort; GS = Generation Scotland. Cohorts with prefix GSE are from the GEO database.
<sup>2</sup> The number of samples in each cohort. Some samples in LBC were measured from the same individual but at different chronological age.

### Estimation of variation in age from using all probes

We used the unrelated individuals from our two largest datasets (GS, N = 2,586, SGPD, N = 1,299) to estimate the proportion of the observed variation in age that is explained when fitting all probes simultaneously, using a mixed linear model analogous to estimating heritability from SNP data^20^. The proportion of variance of age explained by DNA methylation was close to 1 in both cohorts (proportion explained = 1, SE = 0.0036, REML analysis using the software package OSCA^21^ in GS, and 0.99 in SGPD, SE = 0.058), indicating a perfect age predictor can in principle be developed based on DNA methylation data if all probe associations are estimated without error. To demonstrate that this result is not caused by a violation of assumptions, we undertook a permutation test using the same cohorts. We shuffled the ages across individuals and found that DNA methylation did not explain any significant amount of variation in GS (proportion explained = 0, SE = 0.0030) and SGPD (proportion explained = 0.0079, SE = 0.013).

### Building multiple age predictors

To build our age predictors, we collected 14 cohorts and used a common set of 319,607 probes that passed quality control **(Material and Methods)** in all cohorts. We randomly combined 1 to 13 cohorts as a training set, and used the remaining cohorts as test sets. We repeated this step 65 times to generate different training sets with various sample sizes and age spectrum **(Material and Methods)**. We implemented two estimates to evaluate the performance of our age predictors: (1) correlation between predicted age and chronological age in the test data set; (2) Root Mean Square Error (RMSE) of the predicted age in the test data set. Correlation indicates the strength of a linear relationship between the predicted age and chronological age and RMSE reveals the variation of the difference between predicted and chronological age. Two methods, namely Elastic Net^18^ and BLUP^19^ were used. Elastic Net was previously used by Horvath^7^ and Hannum et al^6^. to build their age predictors and BLUP was used to predict age in Peters et al.^22^. These methods differ in how they select probes that are associated with age and how their effects are estimated. Results show that both methods have a decrease of RMSE **(Figure 1)** and an increase of correlation **(Supplementary Figure 1)** when the training sample size increased. The smallest RMSE based on Elastic Net was 2.04 years. This method gave better results with RMSE relative to BLUP for small training sample size, although the difference with BLUP became smaller with increased sample size **(Supplementary Figure 2)**.

**Figure 1:**
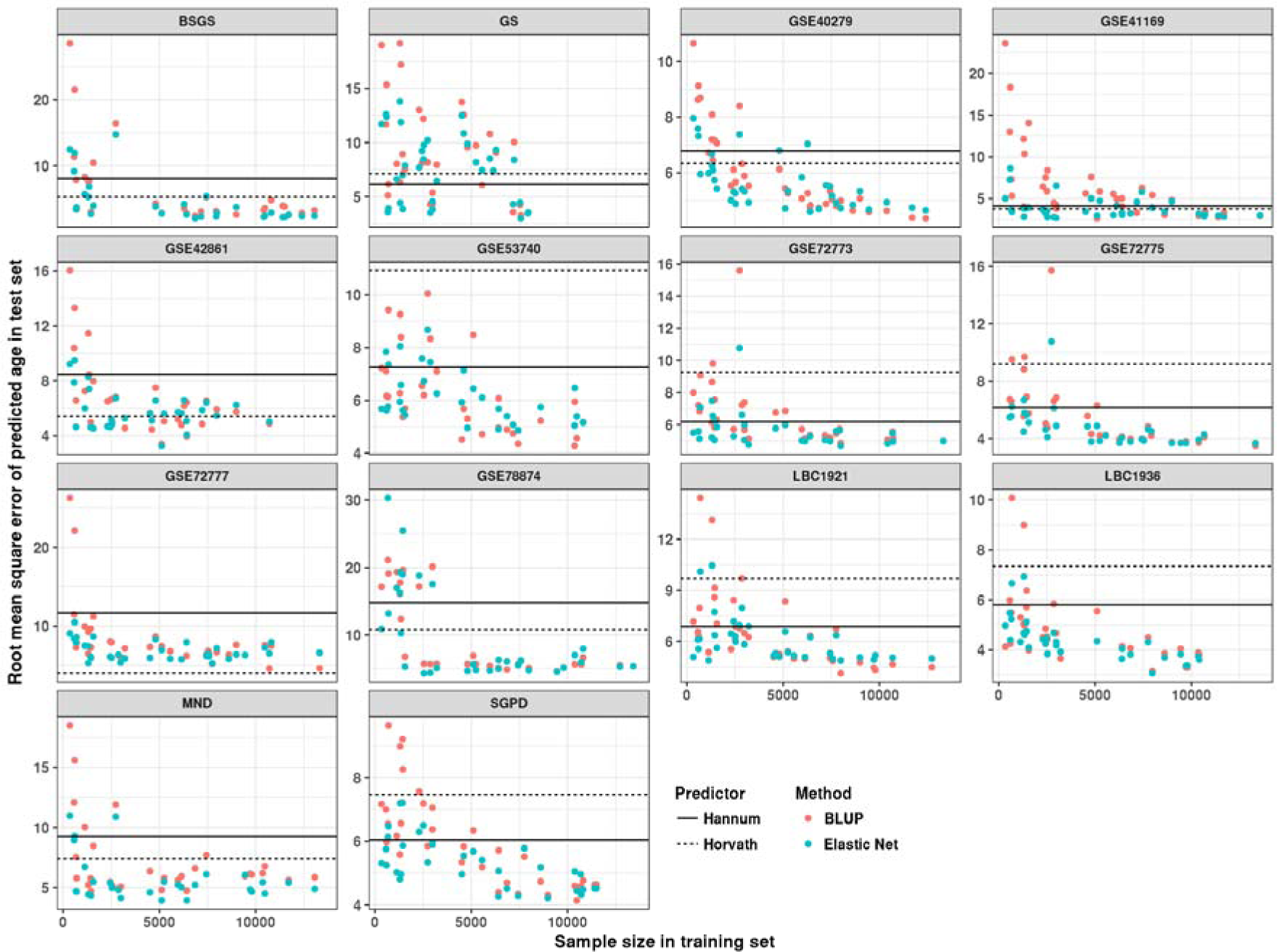
The relationship between training sample size and predictor error measured at the square root of the mean squared error (RMSE) in test data sets. Each point represents the RMSE of the test result based on predictors with different sample size and methods. Prediction results from Horvath are marked as black dash line, and black solid line represents prediction result from Hannum’s age predictor.

### Prediction accuracy and biological ageing

The difference between predicted age from the Hannum/Horvath predictors andchronological age(AAR) was found to be associated with all-cause mortality in later life^13^. To investigate the relationship between the significance of this association and the prediction accuracy of the predictor, we examined the association between AAR and mortality using the updated datain Marioni et al.^13^. These data were from two cohorts: LBC1921 (wave one, N = 436, N_deaths_ = 386) and LBC1936 (wave one, N = 906, N_deaths_ = 214) **(Materials and Methods)**. Age predictors excluding LBC1921/LBC1936 as part of the training set (sample size ranges from 335 to 12,710) were used. We observed a decrease of the test statistics for the effect of AAR on mortality from the survival analysis (Cox regression) with increasing sample size in training data set **(Figure 2)**. No significant (P < 0.05) associations between AAR and mortality was found based on the largest training sample size in either LBC1921 or LBC1936 using BLUP or Elastic Net **(Table 2 and Supplementary Table 1)**. In contrast, results based on the age predictors of Hannum and Horvath were significant (P < 0.05, **Table 2**).

**Figure 2:**
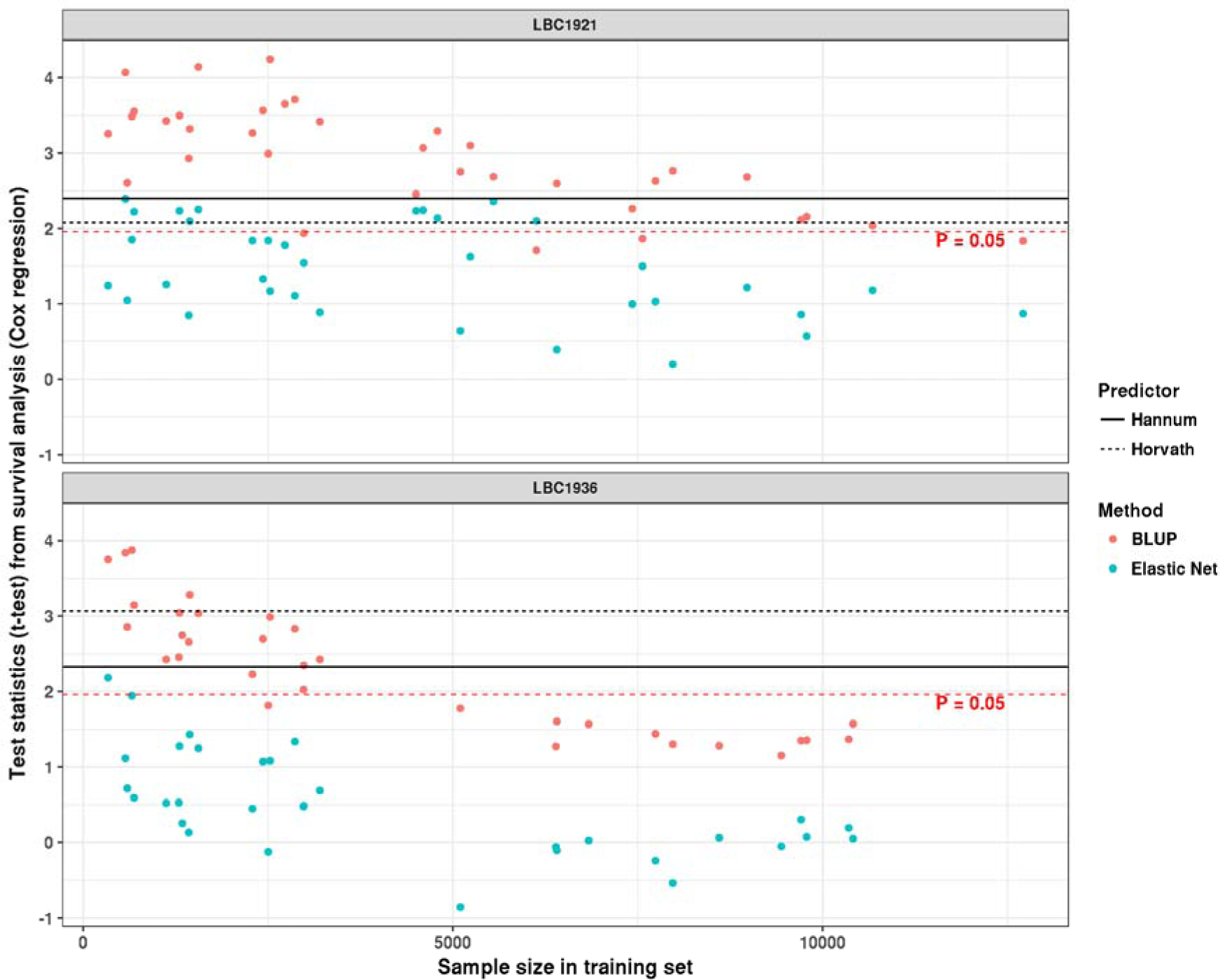
Relationship between the training sample size and the test statistics (t-test) from the association between age acceleration residual (AAR) and mortality. Each point represents the test statistic from the survival analysis based on the predicted ages from predictors with different training sample sizes.

**Table 2:**
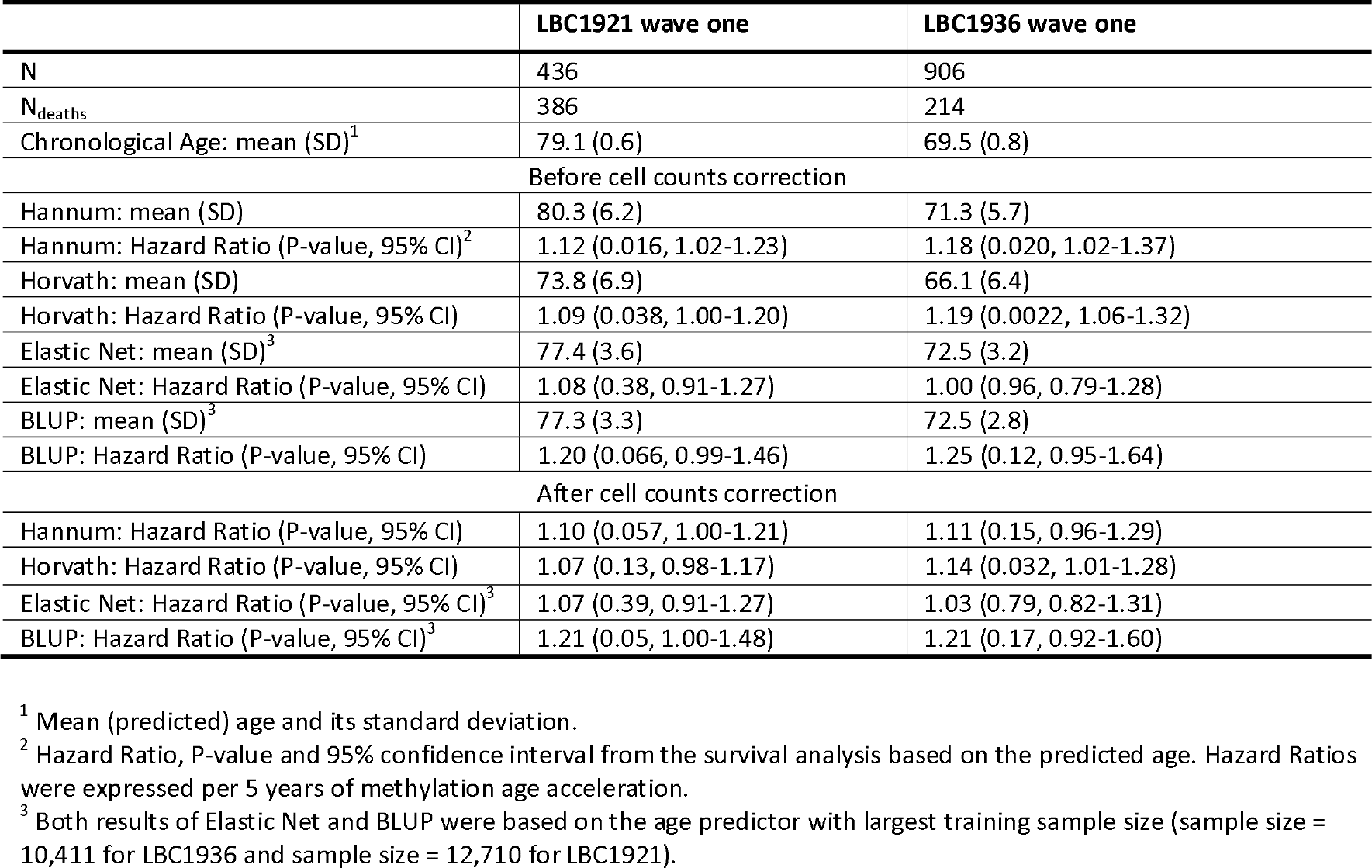
Summary details of two LBC cohorts and the relationship between all-cause mortality and predicted age from different methods (before and after cell counts correction)

### Enrichment analysis on AAR associated probes

Variation in cellular compositions is known to be associated with both DNA methylation^23^ and mortality^24^, which suggests it could be a confounder in the survival analysis. To investigate whether AAR is affected by cellular composition, we applied an epigenome-wide association study (EWAS) on AAR from different predictors. For each predictor, AAR associated (P < 0.05/319,607) probes were selected. We found that the selected CpG sites from age predictors of Hannum and Horvath were enriched in the probes that show heterogeneity in DNA methylation across cell types (cellular heterogeneity probes)^25^ **(Table 3)**, indicating AAR in these predictors was associated with variation in cellular composition. We also observed a decrease of the odds ratio of the enrichment test with the increase of training sample size for both Elastic Net and BLUP based age predictors **(Figure 3)**. No significant enrichment was found for the age predictors based on the largest training sample size **(Table 3)**.

**Table 3:**
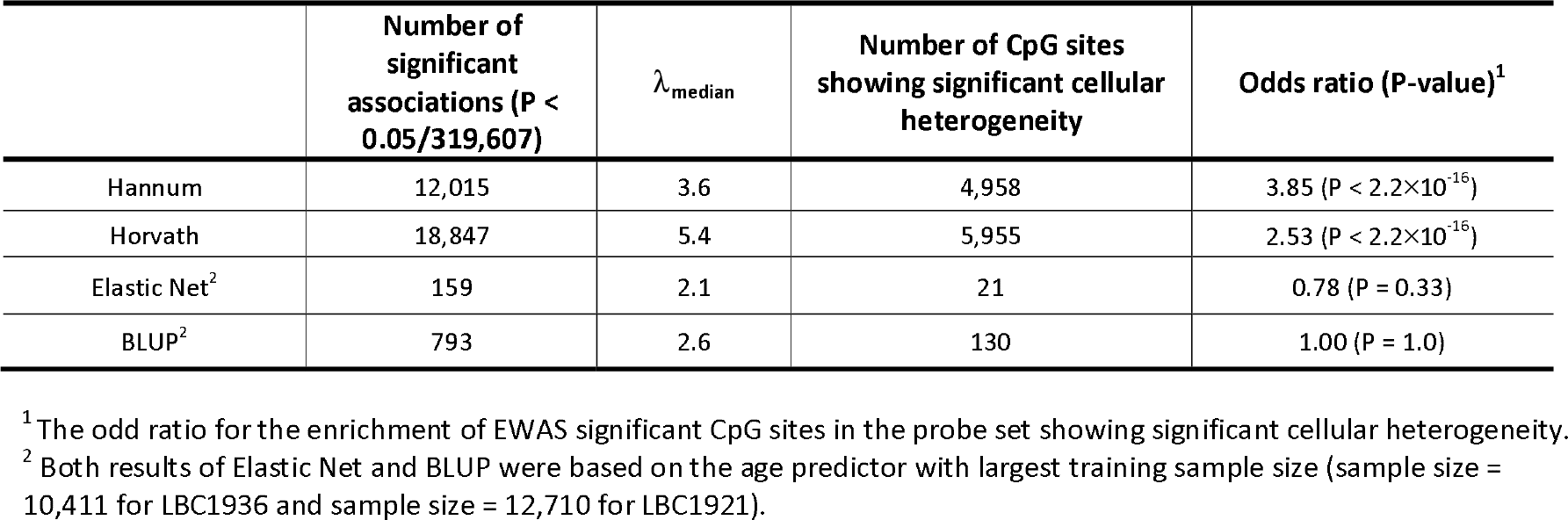
Enrichment test on the AAR associated CpG sites from different methods.

**Figure 3:**
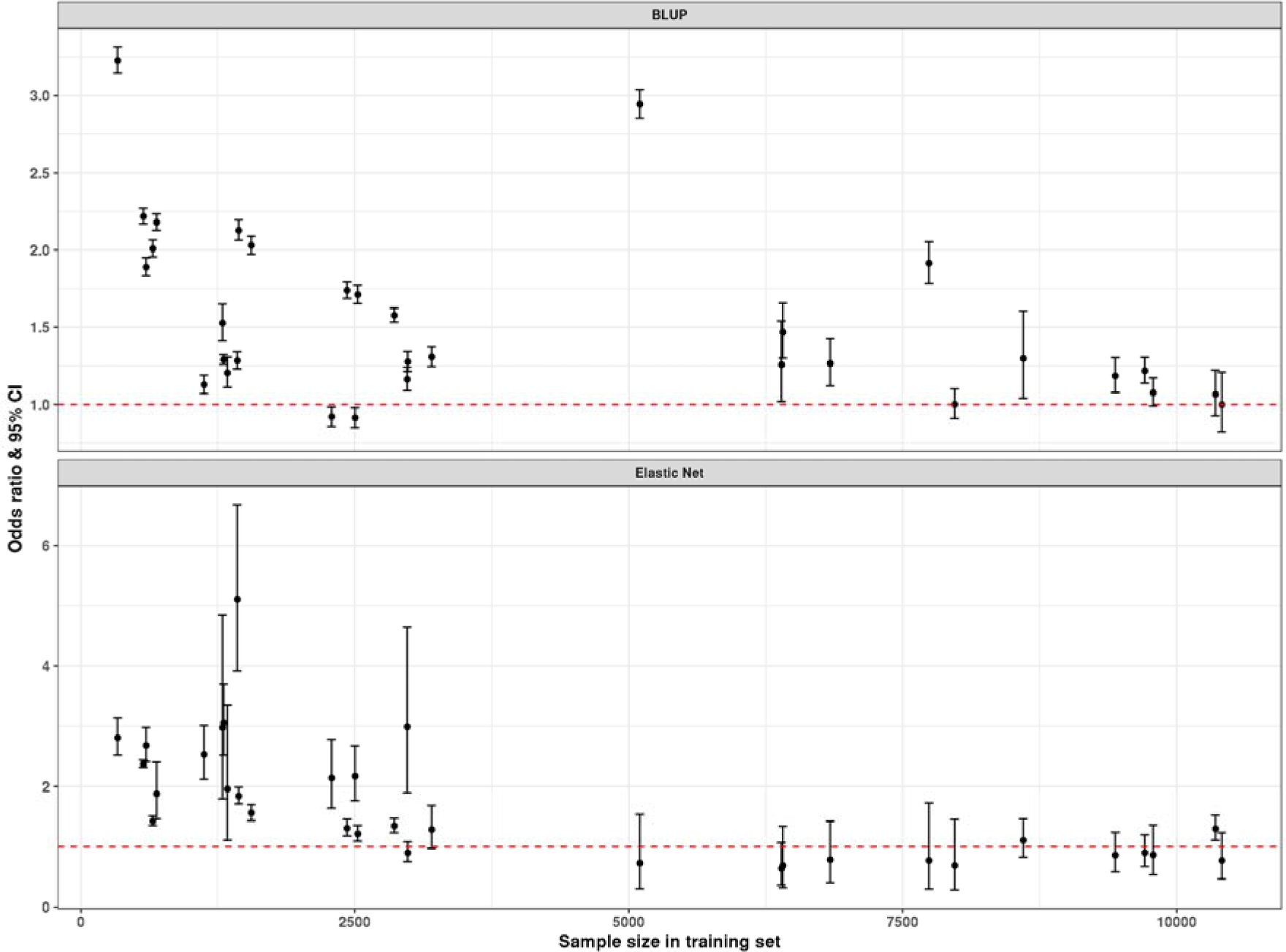
The change of odds ratio from the enrichment test with the increase of training sample size. The enrichment test examines whether AAR associated CpG sites are enriched in probes with cellular heterogeneity.

To examine the effect of variation in cellular content in the association between AAR and mortality, we re-ran the survival analysis based on AAR adjusting for white blood cell (WBC) counts (basophils, eosinophils, monocytes, lymphocytes, and neutrophils) **(Materials and Methods)**. A decrease of the test statistics (from survival analysis) after correcting for the WBC counts was observed, especially when the training sample size is small **(Supplementary Figure 3)**. The effect of AAR on survival is reduced the most (the changes of test statistics from survival analysis are largest) when adding WBC to the models using the Hannum and Horvath age predictors **(Supplementary Figure 3)**. After adjustment for WBC, none of the associations remained significant (P < 0.05) except for the association in LBC1936 based on the predictor of Horvath (P = 0.032). Nevertheless, the significance of this association did not pass the Bonferroni corrected P value threshold (P < 0.05/4) **(Table 2)**. These results suggest that the significant associations between AAR and mortality is biased due to the existence of confounders like WBC counts, and that improved prediction accuracy of the age predictor reduces the effect of these confounders in the survival analysis.

### Other factors related to chronological age prediction

Although the potential effect of confounders limit the “epigenetic clock” to be a biomarker of ageing, a chronological age predictor with good prediction accuracy would be a useful tool in forensics and/or other fields where chronological age is needed but not observed. To determine the factors that explain prediction accuracy, we examined the contribution of age ranges (including absolute age difference between training and test set (Age_diff_) and standard deviation of age (Age_sd_) of the training set) to the RMSE/correlation of the prediction results in the test set by estimating the effect of Age_diff_, Age_sd_ and sample size in the training set on the prediction accuracy jointly **(Material and Methods)**. Results showed that RMSE was significantly associated (P < 0.05) with training sample size in 13 (out of 14) cohorts based on BLUP predictors, confirming that increasing the sample size leads to smaller prediction errors **(Figure 1)**. In addition, eight out of 14 cohorts had a significant (P < 0.05) and positive Age_diff_ effect, indicating similar ages between training and test set results in better prediction accuracy **(Supplementary Table 2)**. Five cohorts were found to have a statistically significant (P < 0.05) Age_sd_ effect on RMSE, suggesting the prediction accuracy benefits from a larger age range of the samples in the training set. Similar results were found based on Elastic Net **(Supplementary Table 3)**. In addition, we did not observe any steady improvement using power-transformed Beta values **(Supplementary Figure 4, Materials and Methods)**, the M values of DNA methylation or the arcsine square root transformed Beta values **(Supplementary Figure 5, Materials and Methods)**.

There is a complex correlation structure in DNA methylation, and the effective number of independent methylation probes was previously reported to be around 200 ^26^, indicating a dense correlation structure. To compare the prediction performance between using the full probe set (319,607 probes) and a pruned probe set (128,405 probes) **(Material and methods)**, we applied the same cross-validation steps to both probe sets using BLUP and Elastic Net. We identified a higher RMSE and a lower correlation for the pruned set **(Supplementary Figure 6)**, indicating a loss of information when using fewer methylation probes for prediction. In addition, we found that probes in the age predictors of Hannum and Horvath were not necessary for age prediction. Compared with these two predictors, better prediction accuracy can still be observed based on the probe set without the probes from these predictors **(Supplementary Figure 7, Material and methods)**, consistent with widespread correlation among probes.

### Age prediction in non-blood tissues

The majority of our samples are from blood, and we observed a significant improvement in the prediction results for the samples from saliva when more blood samples were included in the training set **(Figure 1, Supplementary Figure 1)**. This increase is expected since samples from saliva were reported to exhibit more than 80% contamination by immune cells^27^. To quantify whether our predictor has a good performance in non-blood tissues, we downloaded 13 data sets **(Supplementary Table 4)** that contain samples from other tissues. We compared the performance of our predictor (samples are from blood and saliva, age predictor generated by Elastic Net) with Horvath’s age predictor (samples from multiple tissues, using Elastic Net) in these cohorts and found that our predictor has better performance in samples from endometrium and saliva. On the other hand, Horvath’s age predictor outperformed our predictor in samples from brain (Figure 4). Their performance in other tissues (breast, liver, adipose and muscle) were similar, even though training samples in our predictor are not from these tissues. These results demonstrate that our predictor can also be used to predict the chronological age of samples from non-blood tissues.

**Figure 4:**
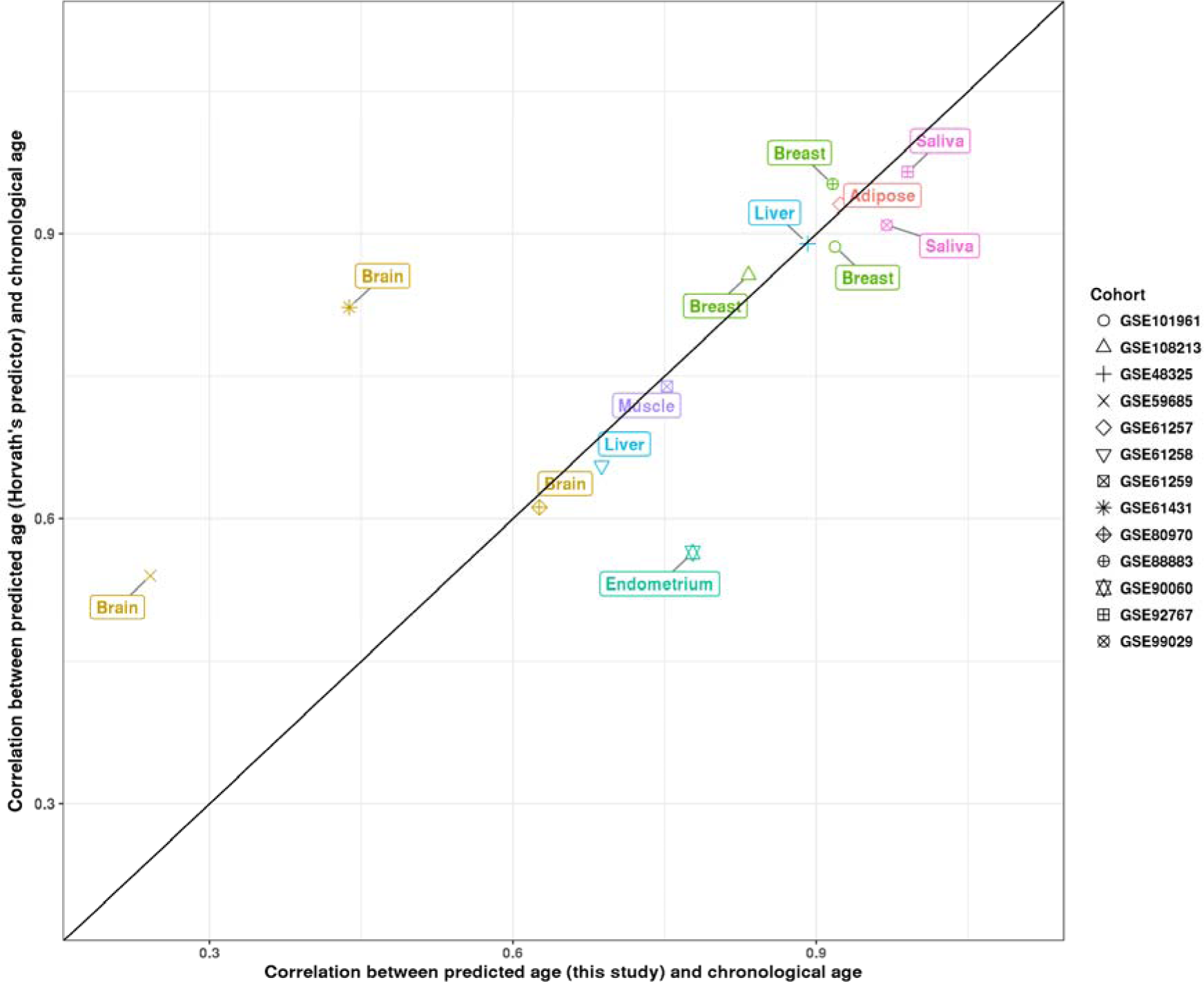
Comparison of prediction performance (correlation) between the predictor from this study (based on Elastic Net) and Horvath’s age predictor in non-blood samples.

## Discussion

We investigated the relationship between the prediction accuracy of an age predictor and its application as a biomarker of ageing. Age predictors with various prediction performance were built based on datasets with different sample sizes (ranging from n=335 to 13566). We ran survival analysis (based on age acceleration residuals AAR) using samples from LBC1921 and LBC1936, with AAR calculated using different age predictors. We observed a decrease in the significance of association between mortality and AAR with the improvement of the age predictor. No significant (P < 0.05) associations were found based on the age predictor with the largest training sample size (Table 2), suggesting the improved prediction of chronological age from DNA methylation limits it as a biomarker of ageing.

We found potential effects of confounders in the association between AAR and mortality in the age predictors of Hannum and Horvath. The AAR associated probes from the age predictors of Horvath and Hannum were enriched in CpG sites showing DNA methylation heterogeneity across cell types, suggesting that AAR from these predictors is affected by variation in cellular composition. The further sensitivity analysis confirmed that, although the AAR from these two predictors are associated with mortality in LBC1921 and LBC1936, no significant (P < 0.05/4) association was observed after adjusting for white blood cell counts. This demonstrates that although the Hannum and Horvath age predictors appear to capture differences in biological ageing between people of the same age, these effects are largely driven by differences in cellular makeup of the samples, and limits their usage as a marker of biological ageing.

We also examined the factors that can affect the accuracy of chronological age prediction, including the effect of the training sample size, the age range of the training samples, the number of probes used and the statistical methods utilised. We found a positive association between the training sample size and the prediction accuracy in test sets. Our predictors showed substantially improved prediction accuracy compared to using the estimated coefficients previously reported by Hannum^6^ and Horvath^7^ in blood samples. Most of this improvement appears to come from simply increasing the experimental sample size in the training set. We also found that increased similarity of ages between samples in the training and test data set can improve the prediction accuracy in the test sets **(Supplementary Tables 2 and 3)**. We provide estimated effect sizes on chronological age from the largest training set of 13,566 individuals for both Elastic Net and BLUP in **Supplementary Table 5**.

Notwithstanding the highly correlated pattern of DNA methylation across the genome, we observed a decline of prediction accuracy when using a correlation pruned probe set, so that including more probes in the training model is beneficial, especially when the training sample size is small **(Supplementary Figure 6)**. The improvement of prediction accuracy could be explained by the decrease of noise effect (such as batch effects) of DNA methylation in age prediction since using more probes can reduce the unexpected effects of the noise. It could also be caused by the existence of many probes with a small correlation with age and the cumulative effect of these may be lost when using a pruned set of probes.

Although most of the samples in our age predictor are from blood, it showed good out-of-sample prediction performance in samples from non-blood tissues. Compared with Horvath’s age predictor, we observed larger correlations (between predicted age and chronological age) in samples from saliva and endometrium, but smaller correlations in samples from brain. These smaller correlations are expected since a large proportion (23.4%) of training samples inHorvath’s age predictor are from brain. Moreover, these two predictors have similar performance in other tissues, which implies that our age predictor is also useful in samples from non-blood tissues.

Our results have several implications for the utility of DNA methylation patterns of age as biomarkers of ageing. From the REML analysis on the SGPD and GS cohorts we estimated that almost 100% of variation in chronological age in those samples could be effectively captured by all the DNA methylation probes on the arrays. For prediction, this implies that for a very large training set a near-perfect predictor of chronological age can be built. Our results showing that larger sample sizes lead to more accurate prediction is consistent with this implication. It is clear that DNA methylation measured in blood is associated with environmental exposures such as smoking, sex and BMI^28-30^. In addition, “age acceleration”, the difference between actual age and that predicted from methylation, has been reported to be associated with a number of outcomes, including mortality^13,16^. However, there is currently no good DNA-methylation-based estimator of an individual’s “epigenetic clock” that is free from confounders (e.g., white blood cell counts) and from prediction error caused by other factors (e.g., measurement error). The difference between actual and predicted age contains both a prediction error term based on unknown factors and possible effects of confounders. These confounders could bias the results when using “epigenetic clock” as a biomarker of ageing, for example the association between “age acceleration” and mortality is confounded by the variation in cellular composition.

## Methods

### Data

We collected 14 data cohorts with samples measured on the DNA methylation 450K chips and Illumina EPIC (850K) arrays **(Table 1)**, eight of which were from the public domain and six datasets from the investigators. Details of the BSGS and LBC cohorts can be found in Powell et al.^31^ and Deary et al.^32,33^. GS is a population and family based cohort recruited through the NHS Scotland general practitioner research network^34,35^. The SGPD cohort is from a collaborative research project on systems genomics of Parkinson’s Disease. Similarly, the MND cohort is from a systems genomics study of Motor Neuron Disease in Chinese subjects (see descriptions in Benyamin et al.^36^). For the purpose of this study, age at sample collection was the focus, disease status and ethnicity of individuals were not considered in any cohort. DNA methylation Beta value at each probe was used for analysis.

A total of 319,607 probes (No Pruned Set) passed our quality control and 128,405 probes (Pruned Set) were retained after pruning based upon the pairwise correlation of probes (see next section). To test the performance of age predictors in non-blood tissues, we downloaded 13 cohorts from GEO database with accession ID GSE61431 (brain)^37^, GSE59685 (brain)^38^ GSE80970 (brain), GSE101961 (breast)^39^, GSE108213 (breast), GSE48325 (liver)^40^, GSE61257 (adipose)^41^, GSE61258 (liver)^41^, GSE61259 (breast)^41^, GSE88883 (breast)^42^, GSE90060 (endometrium)^43^, GSE92767 (saliva)^44^, GSE99029 (saliva)^45^

### Quality Control

All the samples were measured on either the Illumina HumanMethylation450 arrays or Illumina EPIC arrays. Probes with call rate less than 0.95 were removed, and probes found to contain SNPs or potentially cross-hybridizing to different locations were excluded from further analysis^46^. After combining all the samples from different cohorts, a set of 319,607 probes remained (called No Pruned set). Pruning was performed by removing one of two probes on the same chromosome when their correlation (R^2^) was higher than 0.2; this resulted in a set of 128,405 probes (called Pruned set). Both sets were used for further analysis. DNA methylation Beta value was standardized by removing the mean value and divided by the standard deviation for each sample.

### Selection of DNA methylation cohorts

We collected 14 different cohorts in total, including a single cohort (GSE78874) measured in saliva rather than blood tissue. Since DNA methylation is sensitive to batch effects, cell type and tissue type^23^, we applied a PCA analysis (using probes from the No Pruned Set) on the samples from these 14 cohorts to assess the presence of any “outlier” cohorts (i.e. cohorts with a low prediction accuracy from the age predictor based on the other cohorts). All the cohorts were closely matched with the exception of GSE78874 and GS **(Supplementary Figure 8)**. Samples in GSE78874 were from saliva instead of blood, and the samples in GS were measured using Illumina EPIC arrays instead of 450K DNA methylation arrays. To investigate if this difference could potentially adversely influence performance in age prediction for these two cohorts, we used a “leave-one-cohort-out” strategy to leave these two cohorts out as the test set separately and built the age predictor based on the remaining cohorts. We found both of them to have good prediction accuracy (GS: R = 0.98, RMSE = 3.52, GSE78874: R = 0.88, RMSE = 5.39), indicating a small difference between these two cohorts and other cohorts in age prediction. We used all cohorts for subsequent analyses.

### Generation of training set

We generated training sets from the 14 cohorts. Each training set has a certain number of cohorts ranging between 1 to 13. For each number, we repeated random sampling five times. In total, 65 (13 × 5) training sets were generated.

### Proportion of variance of chronological age explained by DNA methylation

The GS and SGPD samples were used in estimating the proportion of variance of chronological age explained by DNA methylation. Among the 5,101 samples in the GS cohort, a subset of 2,586 unrelated individuals, with a genetic relationship coefficient below 0.05 and with no shared nuclear family environment were considered for the analysis. 1,299 unrelated (genetic relationship coefficient < 0.05) individuals with available age information in SGPD were selected. Variance of age was estimated by the REML method implemented in OSCA^21^.

### Prediction algorithm

We compared the age prediction performance of two methods, namely Elastic Net and BLUP. Both methods are based on a linear regression:

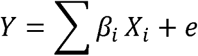

where *Y* is the chronological age, *X_i_* is the DNA methylation of probe *i* and *e* is the Gaussian noise.

Elastic Net is a regularized regression method^18^, and its objective function is defined as:

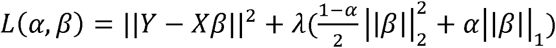

where α and lambda are regularisation parameters. 
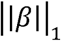
 is defined as 
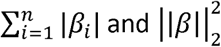
 equals 
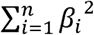
, with *n* the number of probes. *α* is set to 0.5 and is chosen based on cross-validation. We used the implementation of Elastic Net from the Python package glmnet^47^.

BLUP is special case of ridge regression with a fixed *λ*.

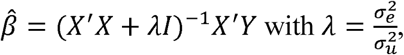

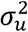
 the variance of the effect size of the probe set, and 
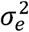
 the variance of the residuals. We used the R package rrBLUP^48^ to build the age predictor, and 
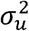
 and 
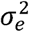
 were estimated using the REML analysis implemented in this package.

### Survival analysis

We followed the same analysis approach as previously described^13^. Briefly, Cox proportional hazards regression models were used to detect the association between the AAR and mortality with age at sample collection and sex as the covariates. AAR is defined as the difference between *m_age_* and chronological age, where *m_age_* is the predicted age correcting for plate, array, position on the array, and hybridisation date (all treated as fixed effect factors), all of which could be confounder in survival analysis. Additional adjustments of AAR were made for WBC counts measured on the same blood samples that were analysed for methylation. Hazard ratios for AAR were expressed per five years of methylation age acceleration **(Table 2)** and per standard deviation of methylation age acceleration **(Supplementary Table 1)**, respectively. Cox models were performed utilizing the ‘survival’ library^49^ in R. Samples from wave one of LBC1921 and LBC1936 were used in this analysis.

### Transformation of DNA methylation

There are non-linear patterns in age-related DNA methylation^50^. To investigate if transformed data can remove the nonlinearity and hence improve the prediction accuracy, we selected eight DNA methylation cohorts with sample size larger than 600 to evaluate the impact of data transformation: LBC1921, LBC1936, GS, BSGS, SGPD, MND, GSE40279 and GSE42861. For each cohort, we randomly selected 70% of the samples as training set, and the remaining 30% were used as test set. Only 50,000 randomly selected probes were used for computational efficiency. Power parameter *λ* (ranges from 0.1 to 2 with 0.05 as the interval) was used to transform the original Beta value of DNA methylation BV to BV^λ^. Only BLUP was used for age prediction because of its low bias. DNA methylation M value and arcsine square root transformed methylation Beta value were also used to compare to raw DNA methylation Beta value in prediction accuracy.

### Age prediction without probes from age predictors of Horvath and Hannum

We compared the probes selected by Elastic Net (based on 13,566 training samples) with those in Horvath’s and Hannum’s age predictors. 11 out of the 514 probes in our analysis were identified in Horvath’s age predictor and 30 in Hannum’s age predictor. In addition, we estimated the squared correlation (R^2^) of DNA methylation between probes selected by Elastic Net and probes from the age predictor of Hannum/Horvath. We found 11 (Elastic Net-Hannum) and 10 (Elastic Net-Horvath) pairs with an R^2^ larger than 0.5 **(Supplementary Figure 9)**, indicating that most of the probes selected by Elastic Net are not strongly correlated with those in the other two predictors. To quantify whether the probes in the Hannum and Horvarth predictors were necessary for age prediction, we re-built our age predictors by excluding these probes. No difference in prediction accuracy was found before and after removing these probes for the BLUP based method **(Supplementary Figure 10)**. The prediction accuracy decreased for the Elastic Net based method; however, its performance was still better than when using the Hannum and Horvath age predictors **(Supplementary Figure 7)**.

## Acknowledgements

This research was supported by the Australian Research Council (DP160102400), the Australian National Health and Medical Research Council (1078037, 1078901, 1103418, 1107258, 1127440 and 1113400), and the Sylvia & Charles Viertel Charitable Foundation. Riccardo Marioni was supported by Alzheimer’s Research UK Major Project Grant [ARUK-PG2017B-10]. Generation Scotland received core support from the Chief Scientist Office of the Scottish Government Health Directorates [CZD/16/6] and the Scottish Funding Council [HR03006]. Genotyping and DNA methylation profiling of the GS:SFHS samples was carried out by the Genetics Core Laboratory at the Wellcome Trust Clinical Research Facility, Edinburgh, Scotland and was funded by the Medical Research Council UK and the Wellcome Trust (Wellcome Trust Strategic Award “STratifying Resilience and Depression Longitudinally” ((STRADL) Reference 104036/Z/14/Z).

## Author contributions

A.F.M and P.M.V conceived and designed the experiments. Q.Z performed all statistical analyses. Q.Z, A.F.M and P.M.V wrote the paper. R.E.M, I.J.D, J.Y and N.W.R advised on statistical methodology, C.L.V, R.M.W, T.L, A.K.H, G.W. M, J.H, D.F, J.F, M.K, T.P, J.P, G.H, J.B. K, I.H, S.L, T.A, P.A.S, G.D.M, S.E.H, P.R, A.D.M, D.J.P, C.S.H, K.L.E, A.M.M, J.G contributed data. All authors read and approved the final manuscript.

